# Heat-induced transposition increases drought tolerance in *Arabidopsis*

**DOI:** 10.1101/2021.11.25.469987

**Authors:** Michael Thieme, Arthur Brêchet, Bettina Keller, Etienne Bucher, Anne C. Roulin

## Abstract

Eukaryotic genomes contain a vast diversity of transposable elements (TEs). Formerly often described as selfish and parasitic DNA sequences, TEs are now recognized as a source of genetic diversity and powerful drivers of evolution. Yet, because their mobility is tightly controlled by the host, studies experimentally assessing how fast TEs may mediate the emergence of adaptive traits are scare. Here, we show that the heat-induced transposition of a low-copy TE increases phenotypic diversity and leads to the emergence of drought-tolerant individuals in *Arabidopsis thaliana*. We exposed high-copy TE lines (hcLines) with up to ∼8 fold increased copy numbers of the heat-responsive *ONSEN* TE (*AtCOPIA78*) to drought as a straightforward and ecologically highly relevant selection pressure. We provide evidence for increased drought tolerance in five out of the 23 tested hcLines and further pinpoint one of the causative mutations to an exonic insertion of *ONSEN* in the *ribose-5-phosphate-isomerase 2* gene. The resulting loss-of-function mutation caused a decreased rate of photosynthesis and water consumption. This is one of the rare empirical examples substantiating the adaptive potential of mobilized stress-responsive TEs in eukaryotes. Our work sheds light on the relationship between TEs and their hosts and demonstrates the importance of TE-mediated loss-of-function mutations in stress adaptation, particularly in the face of global warming.

## Introduction

Plants are constantly exposed to fluctuating environments. To successfully reproduce, they rely on mechanisms that allow them to react and adapt to suboptimal growth conditions. Genetic variation, whether as a result of natural processes or artificially induced, is a prerequisite for adaptation and the evolution of new traits. There is evidence that severe stresses can not only trigger the formation of small-scale mutations (Belfield et al., 2021; Lu et al., 2021) but also increase genetic diversity through the stress-induced activation of transposable elements (TEs) (Negi et al., 2016). Mobile, highly mutagenic (Lisch, 2013; McClintock, 1950) and especially abundant in eukaryotic genomes (Wells and Feschotte, 2020), TEs are believed to facilitate rapid adaptation to challenging environments (Baduel et al., 2021; Li et al., 2018; Naito et al., 2009). Yet, to ensure a limited mutation rate and to safeguard genome stability, TE mobility is usually restricted by epigenetic silencing mechanisms, which in plants involves the RNA-directed DNA methylation (RdDM) pathway (Matzke and Mosher, 2014). As a consequence, only few TE families have been observed transposing *in planta* and therefore the immediate evolutionary consequences of stress-induced TE bursts are largely unknown.

In this context, we study here the functional impact of the *Arabidopsis thaliana* retrotransposon *ONSEN* (AtCOPIA78), one of the best characterized TE-families in plants. Equipped with heat-responsive elements in its Long-Terminal Repeats (LTRs), *ONSEN* can sense the heat stress-response of its host and utilize it to initiate its own lifecycle (Cavrak et al., 2014; Ito et al., 2011; Tittel-Elmer et al., 2010). While the transposition of *ONSEN* is known to create new phenotypes (Ito et al., 2016) and copy numbers have been shown to correlate with the annual temperature range (Quadrana et al., 2016) its adaptive potential has not yet been fully demonstrated. By transiently increasing the rate of naturally occurring transposition events of *ONSEN* through the combination of a chemical inhibition of TE silencing with a heat-shock, we have previously generated genetically stable high-copy TE lines (hcLines) (Thieme et al., 2017) carrying novel insertions of *ONSEN* in wild type (wt) plants of *Arabidopsis thaliana*. Due to climate change and global temperature increase, drought is predicted to constitute one of the most severe environmental constraints to which plants will have to adapt in the near future (Brás et al., 2021; Exposito-Alonso et al., 2019). Here, we used our collection of hcLines to experimentally test whether the heat-induced transposition of *ONSEN* may help individuals to survive in warmer and hence water-limited environments.

## Results and discussion

We grew the S4 generation of 23 hcLines originating from 13 independent heat stressed and transiently de-methylated plants (Thieme et al., 2017) and validated by quantitative PCR (qPCR) an up to ∼8 fold copy number increase (thus up to ∼64 stably inserted *ONSEN* copies; Fig. 1A) compared to Col-0 wt. However, in some hcLines (e.g. hcLine9), no additional *ONSEN* copies were detectable by qPCR even though some were previously validated by sequencing (Roquis et al., 2021). As the hcLines were originally exposed to a combination of heat and the drugs zebularine (Z) and alpha-amanitin (A) (HS+AZ) (Thieme et al., 2017), we controlled for a potential *ONSEN*-independent phenotypic variation caused by epi/genetic changes induced by the heat-stress or the chemical demethylation. To do so, we included a Col-0 wt plant that was propagated on soil and five independent controls, i.e. lines that originated from plants that were also grown *in vitro* but only exposed to control-stress (CS; two lines), heat stress (HS; two lines) or CS plus chemical demethylation (CS+AZ; one line) (Thieme et al., 2017). In accordance with previous observations (Roquis et al., 2021; Thieme et al., 2017), we did not detect an increase in *ONSEN* copy numbers in these control lines when compared to the wt (Fig. 1A).

**Fig. 1.**
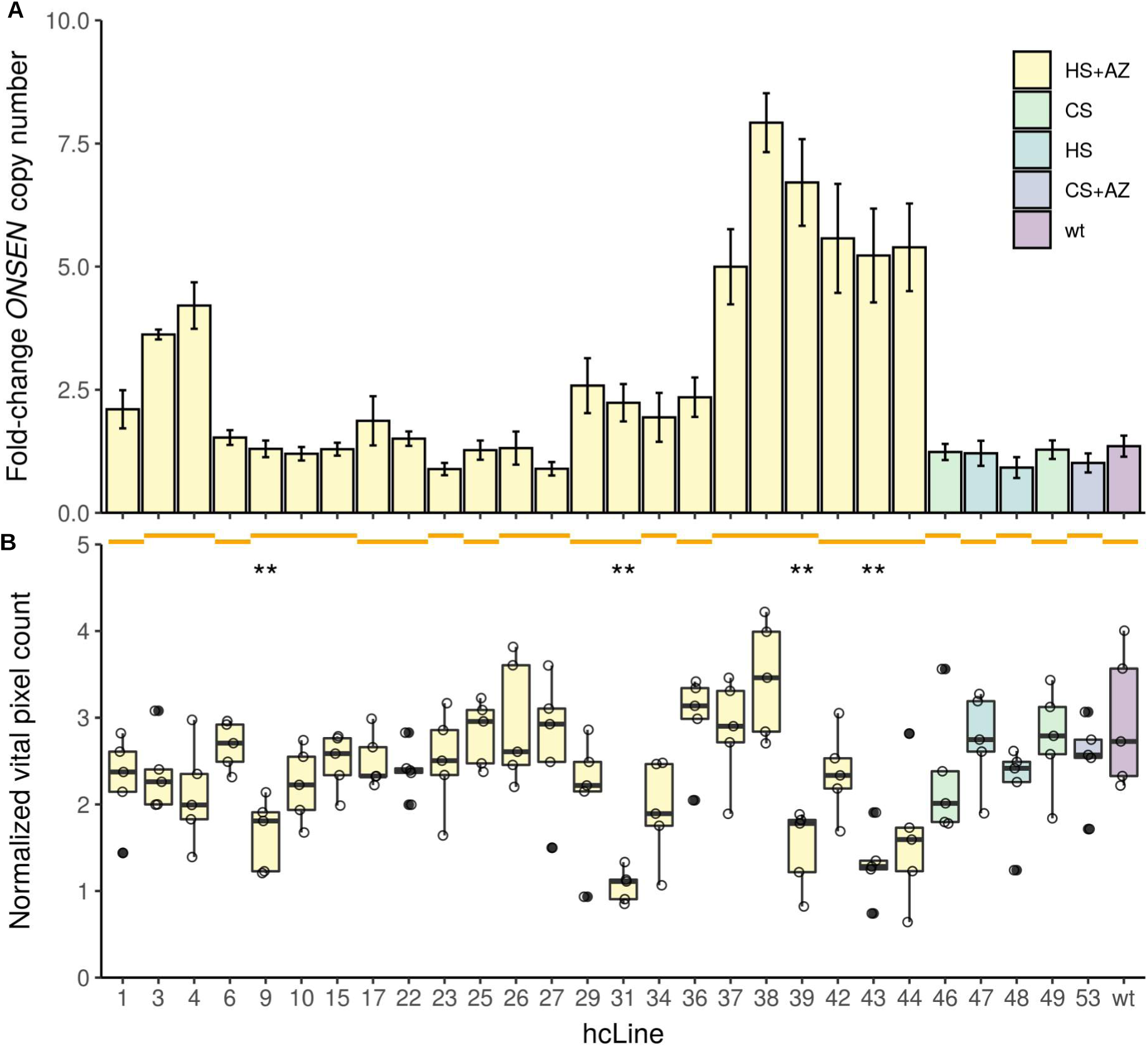
*ONSEN* copy numbers and size of hcLines. Box colors indicate the history of the lines: control-stress (CS), heat-stress (HS), chemical de-methylation (AZ) and wild type propagated on soil (wt). Orange lines spanning multiple hcLines indicate their origin from a common parent. (**A**) Fold-change of *ONSEN* copy numbers measured by qPCR compared to the wt (n=3 technical replicates +/- SD). (**B**) Vital pixel count of hcLines and controls before the occurrence of necrotic leaves (day 8). Significant differences to the wt are indicated. Horizontal line defines median, hinges represent 25th and 75th percentiles, whiskers extend to 1.5 * IQR and outliers are shown as filled dots. n= 5 biological replicates. Wilcoxon-test P< 0.01 (**).

We suspended watering after 36 days of growth and recorded plant development and water loss of the pots every week by taking top view pictures and by weighting the pots. To quantify the growth and the degree of drought-induced leaf senescence, we trained the image-based interactive learning and segmentation toolkit (ilastik) (Berg et al., 2019) to specifically detect living (hereafter vital) and necrotic leaf segments. We first tested the reliability of the prediction by placing one to three punched leaf discs of necrotic or vital segments onto a pot that did not contain a plant. After processing the images with ilastik, we obtained a linear increase of vital and necrotic pixel-counts according to the number of segments placed onto the pots (Fig. S1), confirming the reliability of the method. For the rest of the study, we therefore use vital pixel counts as a proxy for plant size. To assess size variations between the hcLines, we analyzed the pixel counts of predicted vital areas before the appearance of necrotic leaves eight days after watering was suspended (Fig. 1B, Fig. 2A). Although heat stress and the AZ drug treatment have been shown to induce epi/genetic mutations (Belfield et al., 2021; Liu et al., 2015; Roquis et al., 2021), we did not observe differences in growth among our five control lines. Yet, in accordance with the fact that transposition is predominantly associated with fitness loss of the host (Boissinot et al., 2006; Chuong et al., 2017; Roquis et al., 2021; Wilke and Adams, 1992), we found a significantly reduced number of vital pixels compared to the wt for four hcLines (Wilcoxon-test P<0.01, Fig. 1B). Notably, in some cases we also observed a strong size variation between hcLines originating from the same parent (e.g., hcLine37 and hcLine38 vs hcLine39) indicating genetic segregation of the lines (Fig. 1B). We did not find any significant global correlation between *ONSEN* copy numbers and plant size (P>0.05, Fig. S2). Taken together, these observations suggest that single TE insertions rather than the overall *ONSEN* load were responsible for the observed phenotypic variation.

**Fig. 2.**
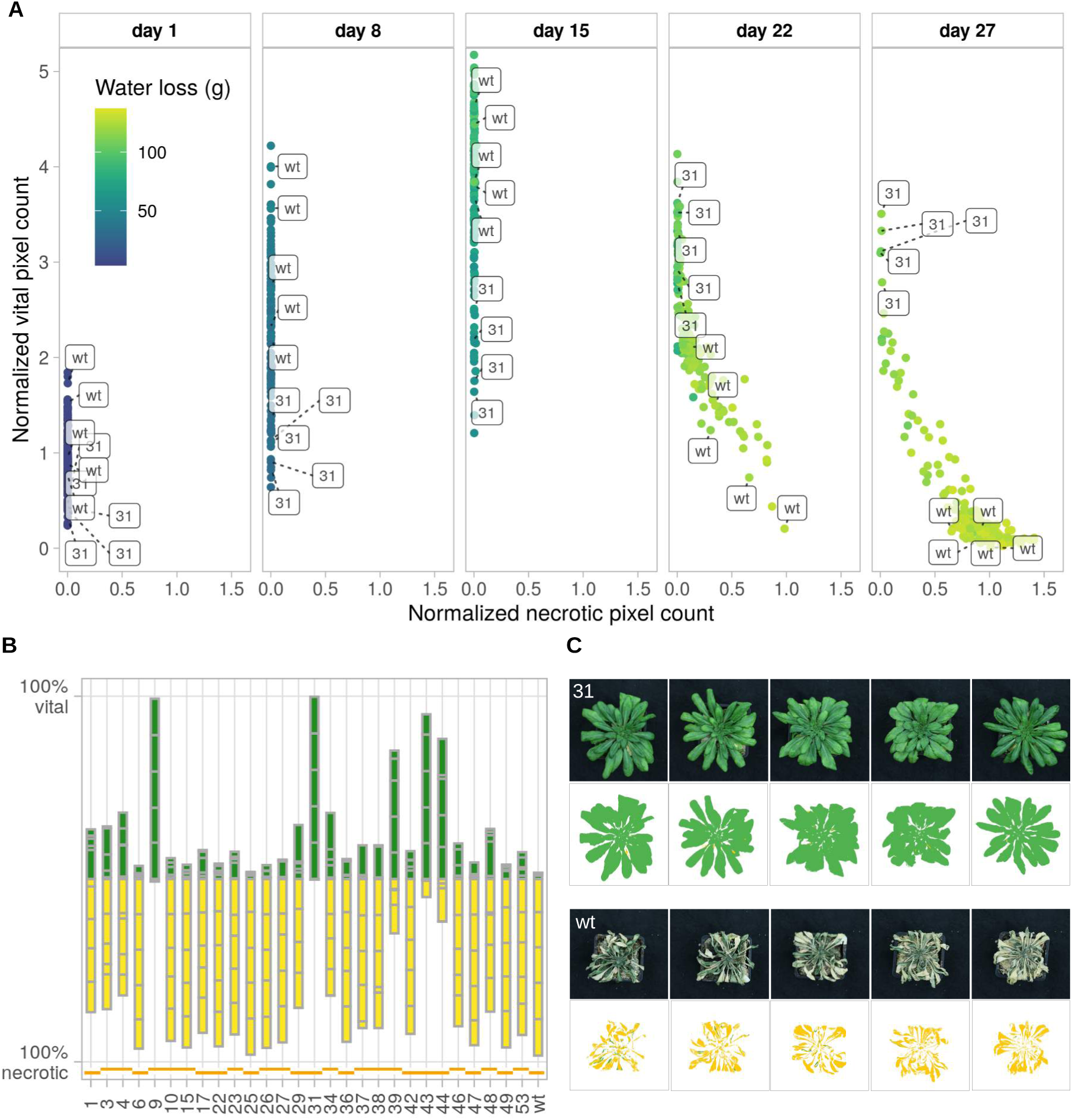
Drought tolerance of *ONSEN* hcLines. (**A**) Pixel counts of living and necrotic tissues during the drought stress. Each dot represents one individual plant and the color code indicates the cumulative water loss over time. n=3-5 biological replicates per line are shown. (**B**) Percentage of vital (green) and necrotic (yellow) tissues of all five replicates per line two days after recovery (day 29). The contribution of each replicate to the total amount of vital and necrotic pixels is indicated with gray bars. Orange lines spanning multiple hcLines indicate their origin from a common parent. (**C**) Original (top) and processed (ilastik, bottom) images of the five replicates of hcLine31 (31) (top) and wt (bottom) on day 29. Predicted vital leaf segments are depicted in green, necrotic segments in yellow.

The machine learning approach allowed us to further track the dynamics of growth and drought-induced necrosis of the individual lines. None of the plants had necrotic leaves until 15 days after watering was suspended, (Fig. 2A). After 27 days, we resumed watering and recorded leaf areas after two days of recovery (day 29). In contrast to the high drought-induced mortality of the five control lines and the wt, five out of the 23 hcLines originating from four independent parental lines were more stress tolerant (mean vital leaf area > 50%) (Fig. 2B). While the wt was most susceptible to drought (0.03 % mean vital area), hcLine31 did not show any signs of necrosis (97.9 % mean vital area, Fig. 2A, B and C). The consistent vitality of all five replicates of hcLine31 indicated that a homozygous mutation was underlying the observed drought tolerance of this line. Because hcLine31 displayed the most drought-tolerant phenotype, we selected it to characterize the functional link between the heat-induced insertion of novel *ONSEN* copies and the observed increase in drought tolerance.

We first used whole-genome re-sequencing data to locate all transposon-insertion polymorphisms (TIPs) of hcLine31 (Roquis et al., 2021). In accordance with its insertion bias towards actively transcribed regions in the *A. thaliana* genome (Quadrana et al., 2019; Roquis et al., 2021), six out of the ten *ONSEN* TIPs detected in hcLine31, were located in exons (Table S1). Notably, we detected a homozygous TIP in the *ribose-5-phosphate-isomerase 2* (*RPI2, At2G01290*), a gene involved in chloroplast photosynthetic capacity (Xiong et al., 2009). Analysis of previously published RNA-seq data of hcLine31 (Roquis et al., 2021) revealed a premature transcriptional stop coinciding with the detected exonic *ONSEN*-insertion in *RPI2* (Fig. 3A).

**Fig. 3.**
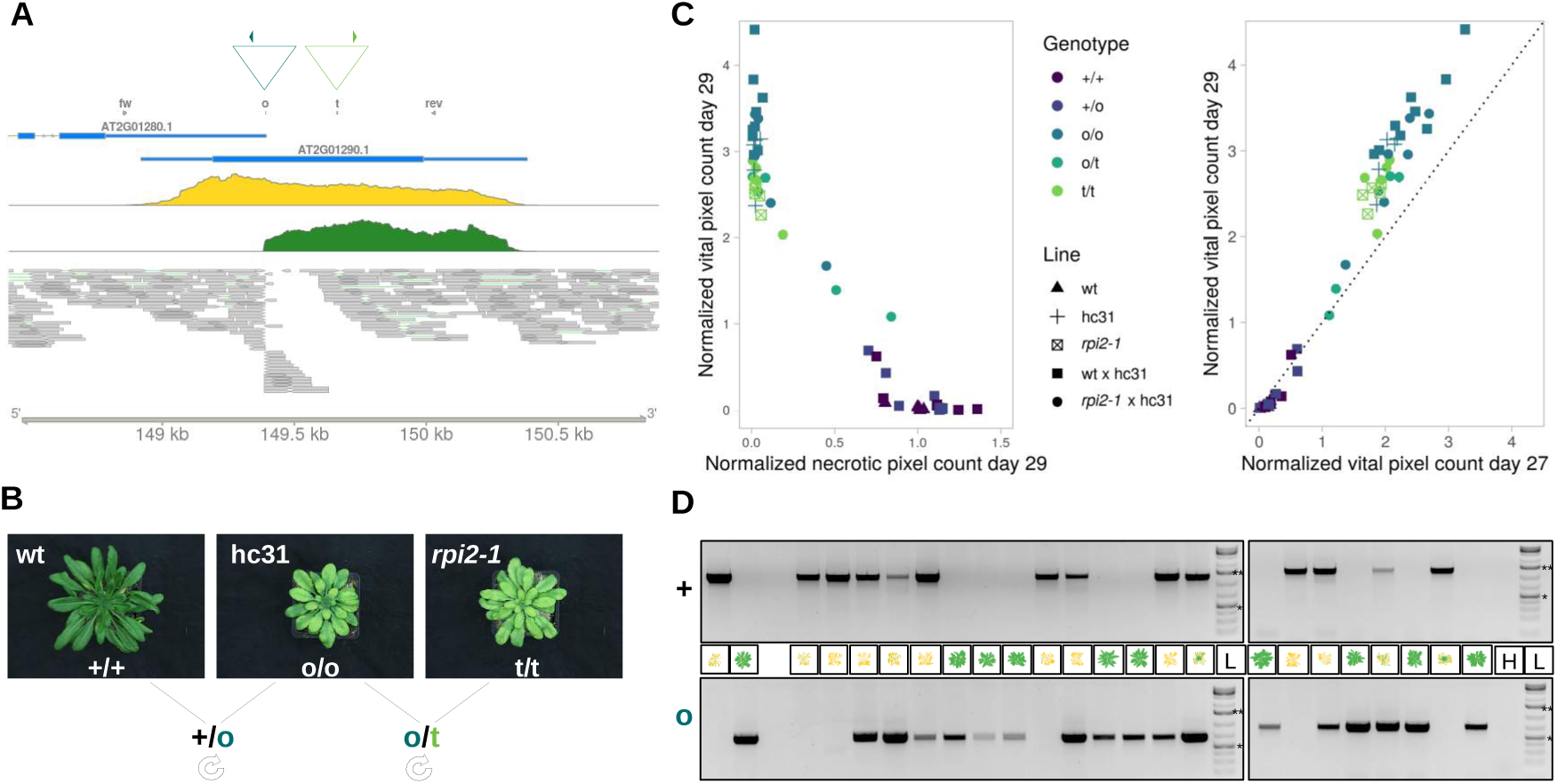
An *ONSEN*-insertion in *RPI2* leads to an increased drought tolerance of hcLine31 (hc31). (**A**) Triangles indicate the insertion site of *ONSEN* (turquoise) and the location of the T-DNA in the *rpi2-1* mutant (green) on chromosome 2. Primer locations used for the genotyping in (**C**) and (**D**) are depicted as filled triangles. Annotation track (blue), RNA-seq coverage of the wt (yellow) and hcLine31 (green) and aligned genomic reads from hcLine31 are shown. (**B**) Images and crossing scheme of the wt, hcLine31, and *rpi2-1*. Genotypes are depicted with + (wt), o (*ONSEN*), and t (T-DNA). **c** Pixel counts of living tissue after two days of recovery (day 29) in relation to necrotic tissues on the same day (left panel) and compared to vital pixel counts before recovery (day 27) (right panel). Shapes indicate the plant line (parental or segregating F2 individuals of the crosses from panel b and colors indicate the genotype of *RPI2*. (D) Geno- and phenotypes (day 29) of the segregating F2 population of hcLine31 x wt. A wt and a hcLine31, (first two lanes) are shown as references. Primers specific for the wt (top gel, +) and the *ONSEN* insertion (bottom gel, o) (see (A)) were used. H= water control, L=GeneRuler 1 kb Plus ladder. 0.5 kb (*) and 1.5 kb (**) bands are marked.

To test whether the *ONSEN* insertion in *RPI2* was the causative mutation for the increased drought tolerance of hcLine31, we used the mutant *rpi2-1* (SALK_022117) as it harbors a homozygous T-DNA insertion in the exon of *RPI2* which also leads to a knock-out of the gene (Xiong et al., 2009). We crossed hcLine31 to the wt and to the *rpi2-1*-mutant (Fig. 3B) and assessed the drought response in the segregating F2 generations obtained from self-fertilization. All F2 individuals that were either homozygous for the *ONSEN* or the T-DNA insertion or that carried both the *ONSEN* and the T-DNA-insertions in *RPI2* survived the drought stress and showed continued growth two days after recovery. In contrast, plants that carried at least one wt allele showed a high degree of drought-induced necrosis and no growth increase two days after recovery from water-limitation (Fig. 3C,D). These results confirmed that the *ONSEN* insertion in *RPI2* results in a recessive loss-of-function mutation causing the increased drought tolerance of hcLine31.

A previous study indicated that the loss of *RPI2* leads to chloroplast dysfunction and reduced chlorophyll content (Xiong et al., 2009). Therefore, we assessed carbon/nitrogen (C/N)-ratios and Soil Plant Analysis Development (SPAD) values, which are directly linked to photosynthetic activity (Ling et al., 2011; Otori et al., 2017), following growth under well-watered conditions and before the emergence of necrotic leaves. SPAD and C/N ratios were significantly reduced (Wilcoxon-test, P<0.05) in hcLine31 compared to the wt (Fig. 4A). As noted earlier, the onset of water-limitation symptoms in hcLine31 occurred significantly later than that in the wt (Fig. 2A), therefore we looked for a general link between plant size and drought tolerance. We first fitted a linear model where water loss (before the occurrence of necrotic leaves at day 8 of the experiment) was entered as the response variable and the plant lines, vital pixel count and their interaction as explanatory variables. While the overall model was significant and explained a large part of the variance (R^2^ =0.72, P < 2.2e-16), we did not detect a significant contribution of the plant line nor of the interaction between the line and vital pixel count, indicating that all hcLines, control lines and the wt had a similar efficiency regarding water use. However, we found that vital pixel count had a significant effect on water loss (P = 5.19e-11).

**Fig. 4.**
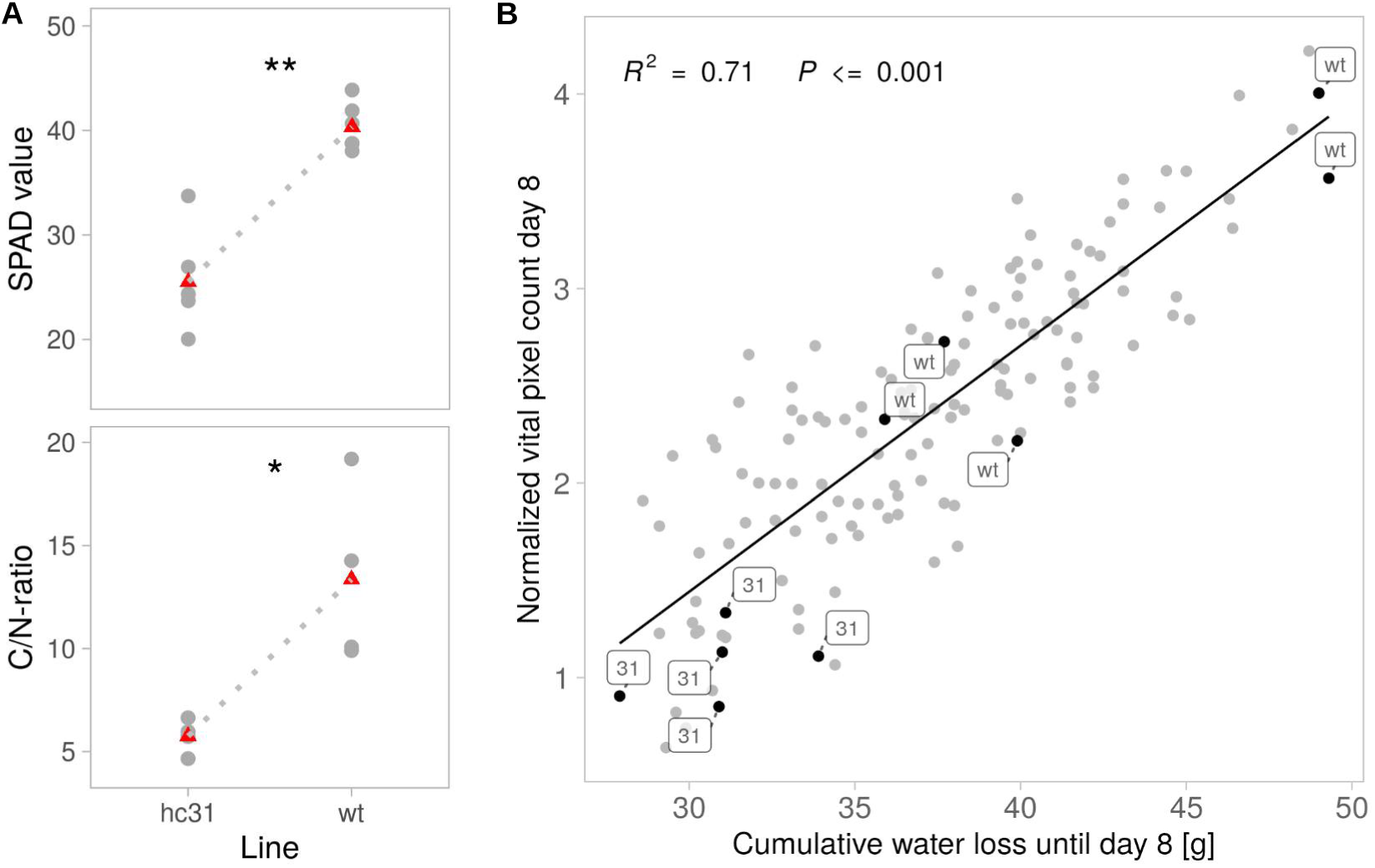
Physiological differences between hcLine31 and wt plants. (**A**) SPAD meter values (top) and C/N-ratio (bottom) of the two lines. n= 4-6 biological replicates. Means are marked with red triangles, Wilcoxon-test P<0.05 (*), 0.01 (**). (**B**) Relationship between pixel counts of living tissues and cumulative water loss until day 8 before the occurrence of necrotic leaves. Linear regression model, adjusted R-squared: 0.71. P<0.001.

This was further confirmed by a reduced model where the line and interaction effects were dropped (Fig. 4B, R^2^ 0.71, P<0.001). These experiments suggested that the *ONSEN* insertion in *RPI2* resulted in reduced photosynthetic capacity leading to slower growth and a reduced water consumption, allowing hcLine31 to escape severe drought stress. Accordingly, we also found that *RPI2* alleles in natural *A. thaliana* ecotypes were quantitatively associated with aridity levels in their local habitats (4.3% of the variance explained; Fig. S3) and concluded that our candidate *ONSEN* insertion in hcLine31 boosts the native function of *RPI2* in respect of adaptation to drought.

Such an insertion of *ONSEN* or other heat-induced TEs could thus provide a selective advantage in the face of global warming (Baduel et al., 2021; Ito et al., 2016; Quadrana et al., 2019). Indeed a correlation between *ONSEN* copy numbers and annual temperature range in natural populations has been reported previously (Quadrana et al., 2016). Yet, because the knock-out of *RPI2* leads to a reduced photosynthetic activity, one may expect large-scale mutations providing such a growth penalty under control conditions to be purged by natural selection. In fact, we do did find any natural TE insertions in *RPI2* (Baduel et al., 2021). This could suggest that such distinct, TE-mediated adaptive effects might be transient and thus difficult to capture with population genomics data. On the other hand, positively selected loss-of-function mutations leading to semi-dwarfness have been reported earlier in *A. thaliana* (Barboza et al., 2013) and the absence of natural TE insertions in *RPI2* could also indicate that this transposition event did not take place in the wild or only occurred in marginal populations not yet sampled. Indeed, TE-mediated mechanisms leading to large-effect mutations might be especially important in less-adapted, frequently stressed populations where drastically altered phenotypes could be advantageous. In agreement with this hypothesis, Baduel et al. (2021) recently pointed towards a link between positive selection of weak alleles of the largest subunit of RNA polymerase V, a key component of RNA-directed DNA methylation, and a globally relaxed silencing of TEs in *A. thaliana* ecotypes growing under extreme conditions.

In the context of climate change, assessing how fast plants will adapt to longer periods of severe heat and drought stress is crucial both for conservation biology and food security (Brás et al., 2021; Exposito-Alonso et al., 2019; Loarie et al., 2009). Selection from standing variation, as opposed to the emergence of new mutations (Hermisson and Pennings, 2017), is classically expected to lead to faster evolution (Barrett and Schluter, 2008). TEs however, may challenge this prediction due to the combined effect of their stress-inducible activity and large mutagenic properties (Baduel et al., 2021). Here, we experimentally confirmed with a real-time setup that a single novel insertion of *ONSEN* can indeed rapidly lead to an selective advantage upon severe water limitation. In *A. thaliana*, several studies have established a functional link between loss-of-function alleles and adaptive traits (e.g. (Gujas et al., 2012; Johanson et al., 2000; Kroymann et al., 2003)) also in the context of adaptation to drought (Monroe et al., 2018; Xu et al., 2019). Indeed, variations in the growth scaling of natural accessions of *A. thaliana* have been shown to be linked to abiotic parameters such as temperature and precipitation (Vasseur et al., 2018). Together with this large body of evidence, our study substantiates the “less is more” hypothesis (Olson, 1999) i.e., that in contrast to intuitive expectations, gene loss may fuel evolution. Whether the observed high frequency of five independent drought-tolerant hcLines could be explained by the insertion preference of *ONSEN* towards actively transcribed H2A.Z-rich regions (Quadrana et al., 2019; Roquis et al., 2021), remains to be elucidated. Selection experiments with populations of hcLines whose TE-composition has not yet been shaped by natural selection will further allow us to extrapolate the overall gain or loss of plant fitness following a heat-induced transposition. In conclusion, our study demonstrates that stress-induced TE mobility can lead to an increased stress tolerance of the host. Because TE activity is family dependent and can be triggered by various abiotic and biotic stresses (Negi et al., 2016), these findings also confirm that TEs, by rapidly modulating gene expression and traits, may provide a powerful mean for plants to keep pace with rapidly changing environmental conditions.

## Materials and methods

### Plant material

*ONSEN* hcLines were generated by treating *Arabidopsis thaliana* Col-0 plants with a combination of a heat-shock and drugs that inhibit TE-silencing, as described previously (Thieme et al., 2017). Briefly, Col-0 seeds were germinated and grown under long-day conditions (16h light) at 24 °C (day) 22 °C (night) on ½ MS medium with 1% sucrose and 0.5% Phytagel, pH 5.8. To reduce TE silencing and increase the rate of *ONSEN* transposition, seedlings were grown analogously on ½ MS medium supplied with a combination sterile filtered zebularine (Z, 40 µM) and ɑ-amanitin (A, 5 mg/ml). After seven days of growth on control ½ MS or medium supplied with A and Z, seedlings were ether exposed to control stress (CS, 24 h at 6 °C followed by 24 h at normal conditions) or heat stress (HS, 24 h at 6°C followed by 24h at 37 °C), then transferred to soil and selfed to obtain the S1 generation. Individual S1 plants originating from plants that were either only exposed to CS or HS or additionally treated with A and Z (AZ) were separated and repeatedly self-fertilized until we obtained the S4 generation. In this study we used 23 *ONSEN* hc-lines originating from 13 plants that were treated with HS+AZ, five independent control lines that were either only exposed to CS (two lines), HS (two lines) or CS+AZ (one line) and the Col-0 wild type that was propagated on soil. The *rpi2-1* mutant (SALK_022117) (Xiong et al., 2009) was obtained from the Nottingham Arabidopsis Stock Centre (Alonso et al., 2003).

### qPCR for *ONSEN* copy numbers

To determine the average *ONSEN* copy numbers of the hcLines and controls used in this study, we extracted DNA of the aboveground parts of at least 24 pooled individuals per line of the S4 generation grown for eight days under sterile conditions on ½ MS medium (1% sucrose, 0.5% Phytagel, pH 5.8) under long day condtions (16h light) at 24 °C (day) 22 °C (night) using the DNeasy Plant Kit (Qiagen). *ONSEN* copy numbers were determined by qPCR using 12 ng total DNA using the KAPA SYBR FAST master mix universal on a C1000 Touch (Bio-Rad) machine. *ACTIN2* (*At*3g18780) was used to normalize DNA levels and DNA of Col-0 served as a control. Three technical replicates were used and data were analyzed with the Bio-Rad CFX Manager 3.1 software. Sequences of oligos are listed in Data S2.

### Identification and visualization of *ONSEN* and T-DNA insertions

Novel *ONSEN* insertions of hcLine31 were identified and characterized recently (Roquis et al., 2021) by whole-genome sequencing and using the Transposable Insertion Finder v1.6 (Nakagome et al., 2014) and the TAIR10 version of the *A. thaliana* Col-0 reference genome (Berardini et al., 2015). To validate the presence and zygosity of the *ONSEN* and T-DNA insertions in *RPI2* in the segregating F2 populations, we designed primers (mto_007 and mto_067) spanning the predicted insertion sites of *ONSEN* and the T-DNA (based on SIGnAL) and combined them with primers specific to *ONSEN* (mto_196) or the T-DNA (LBb1.3 mto_063) (Fig. 3A, Data S2). For the PCRs we used a standard Taq DNA polymerase (Sigma Aldrich) and limited the elongation time to 90 seconds so that an homozygous insertion of the 5-kb *ONSEN* TE or the T-DNA would prevent the formation of a PCR-product. For the genotyping of the F2 populations, we used DNA of homozygous parental plants (wt, hcLine31 and *rpi2-1* (SALK_022117)).

RNA-seq data of one representative biological replicate of the Col-0 wt and hcLine31 exposed to CS and whole-genome sequencing data of hcLine31 were obtained from and analyzed according to a previous report (Roquis et al., 2021). Genomic reads were mapped to the TAIR10 version of the *A. thaliana* genome using bwa mem (v. 0.7.17-r1188) (Li and Durbin, 2009) with the -M parameter set. The insertion site of *ONSEN* was then visualized using the packages Gviz (v. 1.28.3) (Hahne and Ivanek, 2016), rtracklayer (v. 1.44.4) (Lawrence et al., 2009) and the annotation package TxDb.Athaliana.BioMart.plantsmart28 (v. 3.2.2) (Carlson and Maintainer, 2015) using R (v. 3.6.3) (R Core Team, 2020) in Rstudio (v. 1.1.456) (RStudio Team, 2016).

### Drought assay

To obtain comparable and robust results, we ran one comprehensive drought experiment where we tested the S4 generation of hcLines, the control lines and the segregating F2 generations of crosses between hcLine31 and Col-0, and hcLine31 and *rpi2-1* (SALK_022117) in parallel. We included five replicates for each high-copy and control line and for the parents of the cross between *rpi2-1* and hcLine31, and tested 22 F_2_ individuals of the cross of hcLine31 with the wt and 16 F_2_ individuals of the cross of hcLine31 and *rpi2-1* (SALK_022117). Seeds were sown in pots filled with Einheitserde that was incubated with a solution (75 mg/L) of the insecticide Kohinor (Leu+Gygax AG) and kept at 4°C for three days. After stratification, pots were moved into a Hiros climate chamber (Clitec) set to short-day conditions with 10 h light (LED Valoya Ns12 C75/65, ∼120 µmol*m-2*s-1) at 22°C (day) and 19°C (night), with 60% humidity. After ten days of growth, seedlings were piqued into pots filled with equal amounts of Kohinor-treated soil and grown under well-watered conditions for 36 days. The position of pots was frequently shifted to ensure similar growth conditions. Before watering was suspended, pots were again saturated with water and weighted to obtain the maximal water content. One day later (day 1 of the experiment), top-view pictures were taken with a Canon EOS 70D camera on a tripod at the following settings: 5.6 s shutter opening, 1/60 shutter speed, ISO 200. This procedure was repeated three times at an interval of seven days until day 22 of the experiment. Due to technical issues nine out of 193 images (affecting one to two biological replicates of seven different lines and one biological replicate of the F2 of the cross of hcLine31 and *rpi2-*1) from day 15 are missing in the analysis. On day 27, pots were again weighted, top-view pictures were taken, and drought stress was stopped by filling trays with water and allowing the pots to absorb water over night. After two days of regeneration under well-watered conditions, final pictures of the plants were taken (day 29). To account for different zoom levels during the course of the experiment, we took pictures of a white label that later served as a calibrator to normalize predicted vital and necrotic leaf areas. One day after the last pictures were taken, one leaf of each plant from the segregating F2 populations was sampled for DNA extractions and genotyping. Pots were removed from the climate chamber and dried for 8 weeks at room temperature to obtain the dry weight in order to calculated the water content of each pot. Pictures and weight were determined on two successive days (except for day 27); therefore, we extrapolated the weight measurements to determine the water content on the exact day pictures were taken.

### Machine-learning-based prediction of necrotic and vital leaf areas

We used the pixel classification tool of ilastik (v. 1.3.3post3) (Berg et al., 2019) with all 13 features for color/intensity, edge, texture of sigma 0.3, 0.7 and 10.00 selected. We defined three different pixel classes: “background”, “necrotic” and “vital”. We performed an iterative manual training to gradually improve the accuracy of the prediction and finally used 24 images of plants at different stages of the experiment to train the model. For the disc assay, we combined one to three leaf-discs that were punched from vital and necrotic leaves onto soil of a single pot that did not contain a plant. We took three pictures of each combination. To cover a broad spectrum of possibilities, leaf discs were shuffled and/or moved on the pot between pictures. Hence some discs were photographed multiple times but in different combinations and/or positions. Similar to the model training with living and necrotic tissues, we used five images to train ilastik to detect background and the white labels that served as a scale to normalize pixel counts between different days of the experiment. We then processed and exported all image files in ilastik with the following settings: source, “simple segmentation”; convert to datatype, “floating 32 bit”; format, “tif”, and used the getHistogram function of ImageJ (v. 1.53g, Java 1.8) (Schneider et al., 2012) to extract the pixel counts of the areas predicted by ilastik. Pixel counts of vital and necrotic leaves were then normalized using the predicted areas of the size references for each time point of the drought experiment using R.

### SPAD-value and C/N ratio

We grew S4 generation plants under the same conditions and watering regime as for the drought experiment and used a chlorophyll meter SPAD-502 chlorophyll meter (Ling et al., 2011) to determine the Soil Plant Analysis Development (SPAD) values of three or six leaves of six wt and hcLine31 plants that were grown as triplicates in two independent experiments. SPAD values were measured before the occurrence of necrotic leaves 15 days after watering was suspended.

To determine the carbon/nitrogen ratios of the wt and hcLine31 we grew S3 generation plants under well-watered conditions on Einheitserde under short day conditions in a Sanyo MLR-350 growth chamber with 8 h light at 20 °C (day) and 18 °C (night) for 14 weeks. We sampled one leaf from each of four plants of the wt and hcLine31, inactivated them for 30 seconds in a microwave and dried them for eight days at 60°C. Plant material was then ground for 3 minutes at 30 hz with an oscillating mill (M M 400, Retsch, Germany). Then, 2 mg of plant material were put in tin capsule and C/N-ratios were analyzed with a thermal conductivity detector by the Basel Stable Isotope Lab.

### *RPI2* analysis in natural accessions

We used the vcf file produced by The 1001 Genomes Consortium (Alonso-Blanco et al., 2016) to extract SNPs for 1135 sequenced accessions of *A. thaliana*. To limit the effect of the phylogenetic relationship in further analyses, we used the function – relatedness2 from vcftools (Danecek et al., 2011) to keep only ecotypes with a kinship coefficient k <0.5. For the remaining ecotypes, bioclimatic variables (https://www.worldclim.org/data/worldclim21.html) and Global Aridity Index (https://cgiarcsi.community/data/global-aridity-and-pet-database/) were extracted using the Rpackages raster (v. 3.5-2; (Hijmans and van Etten, 2012)) and rgdal (v. 1.5-27; (Keitt et al., 2010)). We fitted a linear-mixed model with the R package lme4 (v. 1.1-27; (Bates et al., 2015)) to test the association between aridity levels averaged over May to August and SNPs in *RPI2* harboring a minor allele frequency (maf) > 0.3. We added the admixture groups defined by The 1001 Genomes Consortium (Alonso-Blanco et al., 2016) as a random effect to account for population structure. We used QGIS (v. 3.16; https://www.qgis.org/en/site/) to display *RPI2* alleles in Eurasia. For the rest of the manuscript, all statistical analyses were performed in R and parametric or non-parametric tests were used according to the sample size and data distribution.

## Supporting information

Data S1 SNPs in RIP2 and bioclimatic variables for the subset of 596 ecotypes

Data S2 Names and sequences of oligos used for the qPCR to determine ONSEN copy numbers and for genotyping of RPI2.

## Data and materials availability

Raw and segmented images and ilastik classifications were uploaded to figshare. Seeds are available upon request to.

### Supplementary tables and figures

**Table S1.** Novel *ONSEN* insertions in hcLine31. Location, description (Araport11) and zygosity (-/- homozygous *ONSEN*, +/- heterozygous) of predicted *ONSEN* insertion sites adapted from (Roquis et al., 2021).

| *chr* | *coordinates* | *context* | *ID* | *description* | *zygosity* |
| --- | --- | --- | --- | --- | --- |
| 1 | 21761950– 55 | exon | *AT1G58602* | LRR and NB-ARC domains-containing disease resistance protein | (-/-) |
| 2 | 149387- 91 | exon | *AT2G01290* | cytosolic ribose-5-phosphate isomerase | (-/-) |
| 3 | 19300993– 97 | exon | *AT3G52020* | serine carboxypeptidase-like 39 | (+/-) |
| 3 | 22631755- 59 | exon | *AT3G61150* | homeodomain GLABROUS 1; HD-ZIP IV family. | (+/-) |
| 5 | 853776- 80 | exon | *AT5G03435* | Ca2+-dependent plant phosphoribosyltransferase family protein | (-/-) |
| 5 | 10632816– 20 | TE | *AT5G28626* | AT5TE38720; SADHU; Sadhu non-coding retroTE family | (-/-) |
| 5 | 18850327- 31 | promoter | *AT5G46490* | disease resistance protein (TIR-NBS-LRR class) family | (-/-) |
| 5 | 21050239- 43 | intron | *AT5G51800* | protein kinase superfamily protein | (-/-) |
| 5 | 21602030- 34 | exon | *AT5G53240* | hypothetical protein (DUF295) | (+/-) |
| 5 | 22846432- 36 | intron | *AT5G56400* | FBD, F-box, Skp2-like and Leucine Rich Repeat domains containing protein | (+/-) |

**Fig. S1.**
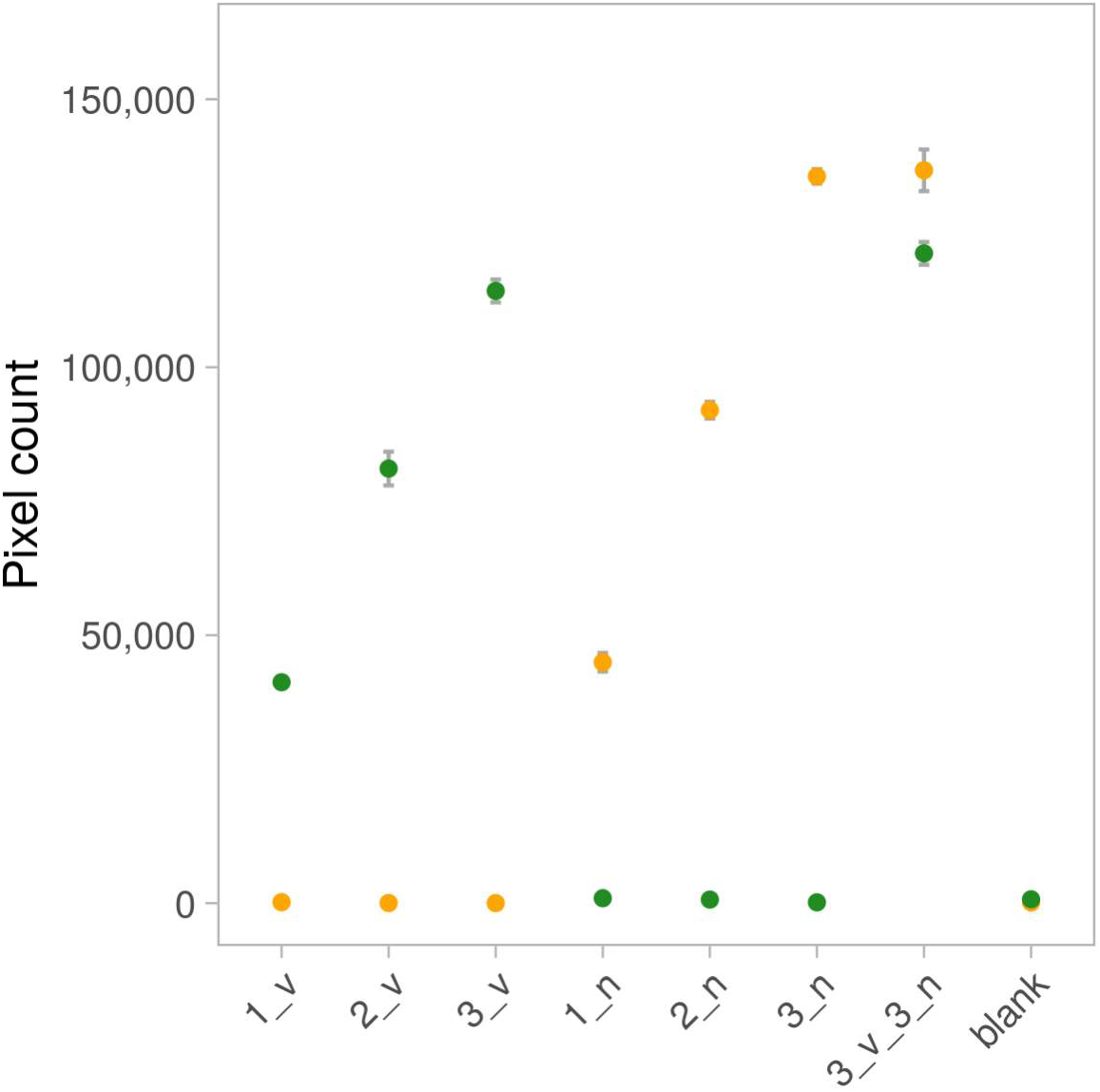
Leaf disc assay to validate the accuracy of the pixel counts for the machine learning-based prediction of vital (green) and necrotic (yellow) segments using ilastik. One, two or three discs were punched from vital (v) or necrotic (n) leaf tissues and placed onto soil of an empty pot (blank). n=3 technical replicates.

**Fig. S2.**
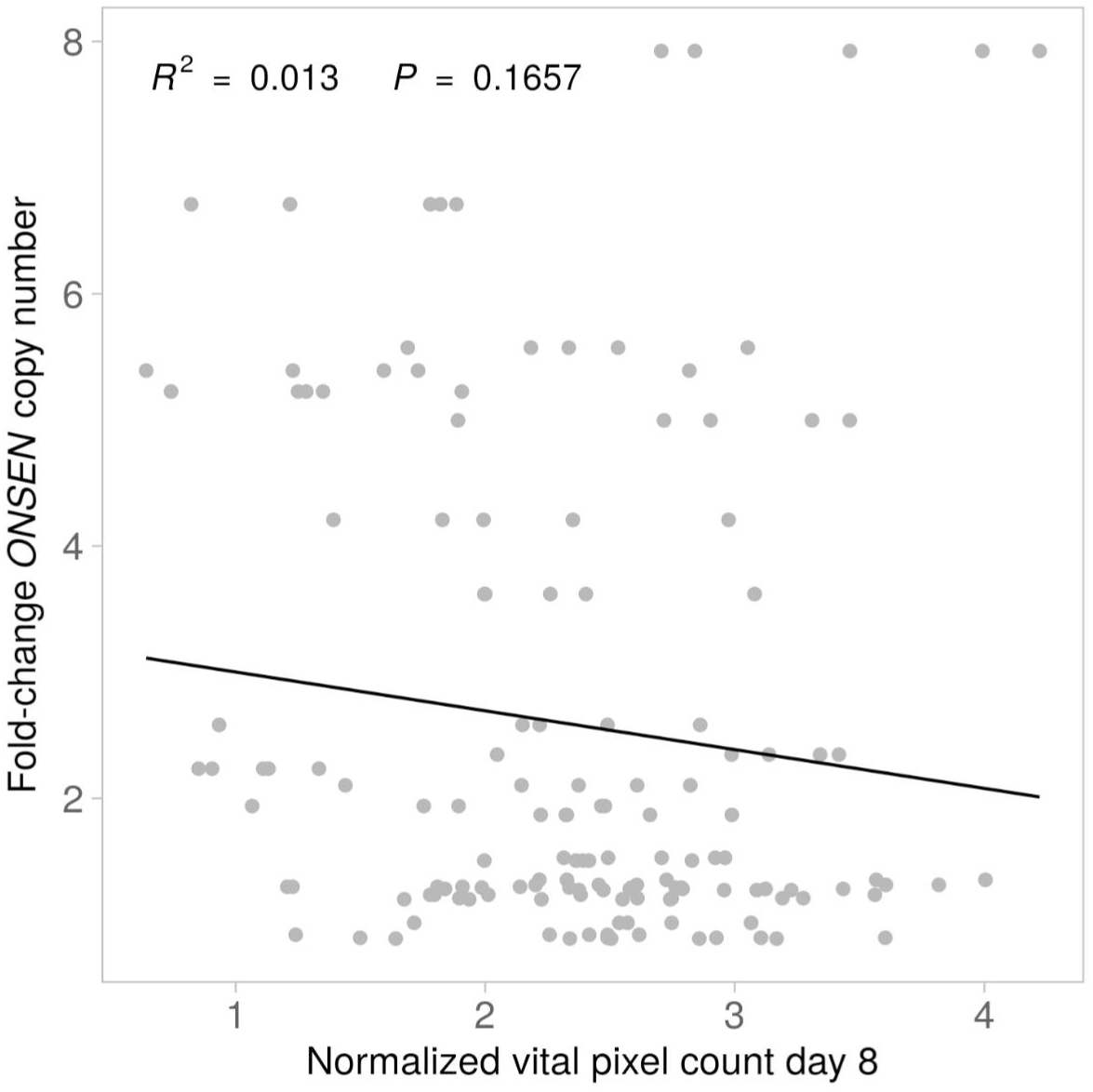
Pixel count of vital tissues in all tested lines before the occurrence of necrotic leaves (day 8) (n=5 biological replicates) and the fold change of *ONSEN* copy numbers measured by qPCR relative to the wt (n=3 technical replicates). Linear regression model, adjusted R-squared: 0.013, P=0.1657.

**Fig. S3.**
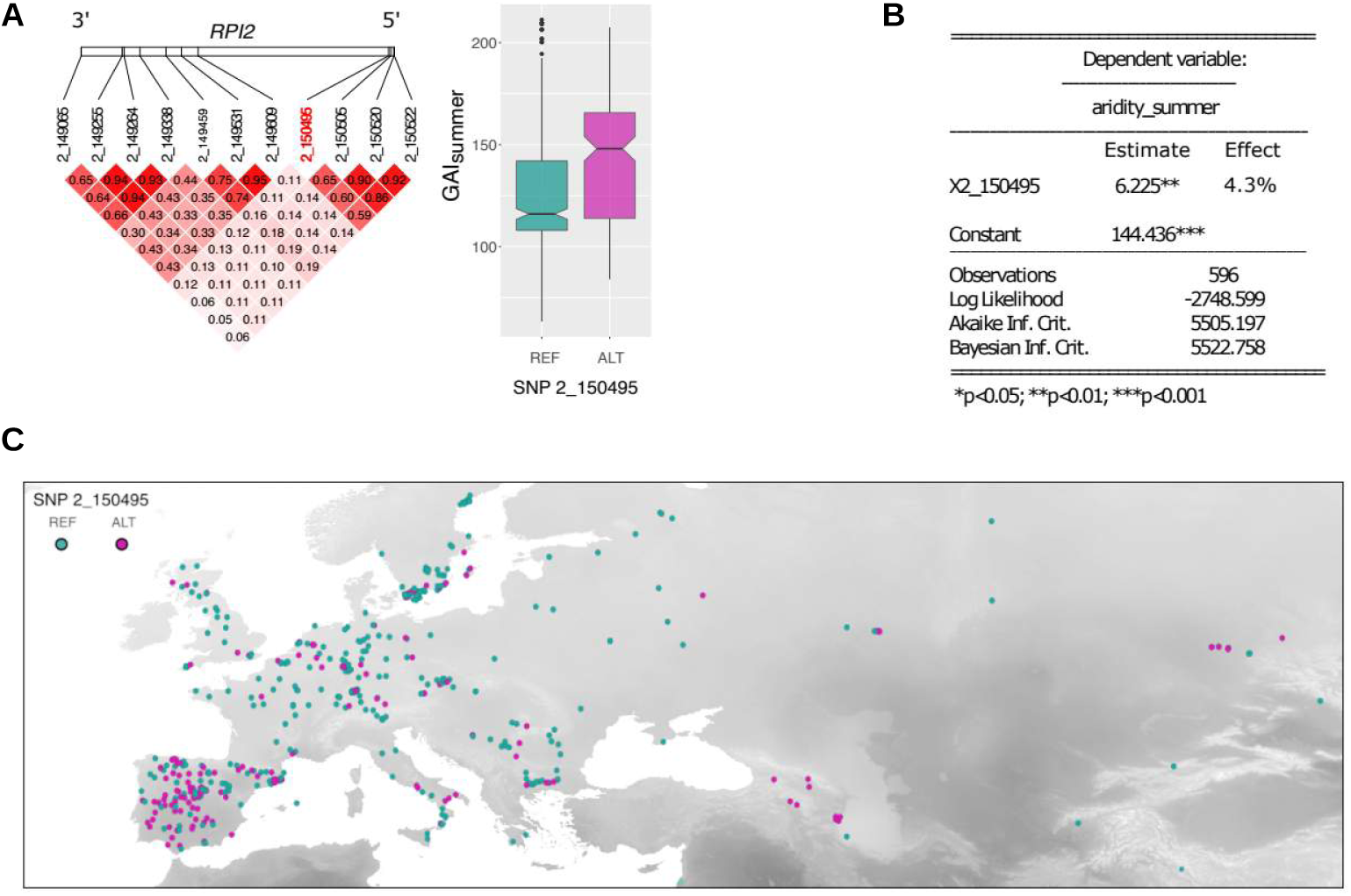
The association between aridity and *RPI2* variations in natural populations of *A. thaliana*. (**A**) Linkage disequilibrium among the 11 SNPs located in *RPI2* (left panel) and the association between SNP 2_150495 and Global Aridity Index in summer (GAIsummer; right panel). (**B**) Linear mixed-model results based on 596 accessions. (**C**) Occurrence of the reference and alternative alleles of SNP 2_150495 in Eurasia. The intensity of the background map displays aridity levels in July from low (light grey) to high (dark grey).

### Supplementary data

**Data S1** SNPs in *RIP2* and bioclimatic variables for the subset of 596 ecotypes.

**Data S2** Names and sequences of oligos used for the qPCR to determine *ONSEN* copy numbers and for genotyping of *RPI2*.

**Data S3** Raw data of pixel counts and weight, SPAD values and C/N-measurements.

## Acknowledgments

We would like to thank Anja Schmutz for her assistance with R, Mahendra Mariadassou for his kind support with statistics and Yann Bourgeois and Christoph Stritt for their comments on the manuscript. We further want to thank Ansgar Kahmen and Thomas Boller for their involvement at the beginning of the project.

## Funding

University of Zurich Research Priority Programs (URPP) *Evolution in Action* (MT, BK, ACR)

European Commission PITN-GA-2013-608422-IDP BRIDGES (MT) Freiwillige Akademische Gesellschaft Basel (MT)

SNSF 31003A_182785 (ACR, BK)

European Research Council (ERC) under the European Union’s Horizon 2020 research and innovation program 725701, BUNGEE (EB)

## Author contributions

Conceptualization: MT

Data acquisition: MT, BK, AB

Data analysis MT, ACR

Funding acquisition: MT, EB, ACR

Writing - original draft: MT

Writing - review & editing; MT, ACR, EB, AB, BK

